# Utilizing the Scale-Invariant Feature Transform Algorithm to Align Distance Matrices Facilitates Systematic Protein Structure Comparison

**DOI:** 10.1101/2023.11.14.566990

**Authors:** Zhengyang Guo, Yang Wang, Guangshuo Ou

## Abstract

Protein structure comparison is pivotal for deriving homological relationships, elucidating protein functions, and understanding evolutionary developments. The burgeoning field of *in-silico* protein structure prediction now yields billions of models with near-experimental accuracy, necessitating sophisticated tools for discerning structural similarities among proteins, particularly when sequence similarity is limited. Here, we have developed the ADAMS pipeline, which synergizes the distance matrix alignment method with the scale-invariant feature transform (SIFT) algorithm, streamlining protein structure comparison on a proteomic scale. Utilizing a computer vision-centric strategy for contrasting disparate distance matrices, ADAMS adeptly alleviates challenges associated with proteins characterized by a high degree of structural flexibility. Our findings indicate that ADAMS achieves a level of performance and accuracy on par with Foldseek, while maintaining similar speed. Crucially, ADAMS overcomes certain limitations of Foldseek in handling structurally flexible proteins, establishing it as an efficacious tool for in-depth protein structure analysis with heightened accuracy.

## Introduction

Currently, sequence similarity searches dominate protein annotation and analysis, aiming to identify analogous sequences that can offer insights into the molecular, cellular, and structural attributes of the subject sequence (Altschul, et al., 1990; Sievers, et al., 2011). Yet, even with the achievements of this approach, it struggles to annotate a myriad of proteins due to the inherent challenges in identifying distant evolutionary connections based solely on sequences. Identifying similarities between protein structures through three-dimensional (3D) superposition provides enhanced sensitivity in recognizing homologous proteins (Liu, et al., 2008). Given the accessibility of high-caliber structures for any given protein, leveraging structure comparison can refine inferences on homology and offer deeper insights into structural, functional, and evolutionary aspects.

Recent advancements in *in-silico* protein structure prediction are propelling the fields of structural biology and bioinformatics forward. Pioneering AI models such as AlphaFold2 (Jumper, et al., 2021), RoseTTAFold (Baek, et al., 2021), and ESMFold (Lin, et al., 2023) have made it possible to generate billions of protein structure models with near-experimental quality, representing a treasure trove of biological insights. The European Bioinformatics Institute currently boasts a repository of over 214 million structures, as predicted by AlphaFold2 (David, et al., 2022), while the ESM Atlas holds an impressive catalog of over 617 million metagenomic structures as deciphered by ESMFold (Lin, et al., 2023). However, this wealth of structural data demands advanced mining tools capable of analyzing, identifying structurally similar proteins, and shedding new light on those with limited sequence similarity. Despite years of dedicated efforts to enhance the accuracy and speed of structural aligners, current mining and comparison tools still have limitation in their accuracy and speed, struggling to manage the burgeoning scale of structure databases.

Traditional protein structure comparison tools such as CE (Combinatorial Extension) (Shindyalov and Bourne, 1998), Dali (Holm, 2020), and TM-align (Zhang and Skolnick, 2005) are foundational in structural bioinformatics. However, their efficiency wanes when handling vast datasets, primarily due to their inherent time complexity. One significant bottleneck is their dependence on rigid-body superposition, which presupposes that the entire protein structure can be optimally aligned as a single entity. This assumption can lead to prolonged iterations to achieve alignment optimization, often resulting in the alignment of one region at the expense of another (van Kempen, et al., 2023). The existence of flexible regions, conformational changes, or significant structural variations can exacerbate this problem, rendering traditional methods time-consuming and unsuitable for large-scale protein structure comparisons.

Foldseek introduces an innovative solution to the conundrum of protein structure alignment through local structure embedding (van Kempen, et al., 2023). This method encodes the structural context of an amino acid into a designated code from a predefined structure code book. Consequently, the amino acid is substituted by its structural descriptor. The ensuing sequence, replete with structural details, is then amenable to standard sequence alignment methodologies (Steinegger and Söding, 2017). The genius of Foldseek lies in its reconceptualization of a structural alignment task as a sequence alignment problem, delivering unparalleled efficiency. This approach also provides a degree of solution to the complex task of simultaneous alignment of divergent protein regions. Nonetheless, it’s pivotal to recognize that while structural embedding offers significant benefits, it may be susceptible to intrinsically disordered regions (IDR) that lack a fixed three-dimensional structure and are often characterized by their flexibility and disorder (Trivedi and Nagarajaram, 2022). Such variances, perceived as noise generated by structure prediction, stem from the inherent flexibility of certain protein conformations (Alderson, et al., 2023). This sensitivity is a noteworthy consideration when deploying Foldseek, especially when the precision of structural representation is paramount.

Several studies have estimated that a significant portion of human proteins contain IDRs. Some estimates suggest that approximately 30-40% of human proteins may contain IDRs to varying degrees (Fig. S1A, B). These IDRs play important roles in various cellular processes, including protein-protein interactions(Hultqvist, et al., 2017), signaling (Bondos, et al., 2022), and regulation (Babu, et al., 2011). Here, we present a novel algorithm, termed <u>a</u>lign <u>d</u>ist<u>a</u>nce <u>m</u>atrix with <u>s</u>cale invariant feature transform, or ADAMS succinctly. Utilizing a computer vision-centric strategy (Lowe, 2004) for contrasting disparate distance matrices, ADAMS adeptly alleviates challenges associated with proteins characterized by a high degree of structural flexibility. In essence, ADAMS operates at a speed comparable to Foldseek, establishing it as a practical choice for comprehensive protein structure comparison with elevated precision.

## Result

### Overview of the ADAMS algorithm

ADAMS integrates the Distance Matrix Alignment (Dali) method (Holm, et al., 2023; Holm and Sander, 1993) with the Scale-Invariant Feature Transform (SIFT) algorithm (Lowe, 2004) (Fig. 1 A). Initially, a distance matrix is derived from the *C*_*α*_ backbone coordinates of a designated PDB structure or a protein structure predicted by AlphaFold2. Following this, keypoints are extracted using the OpenCV-SIFT algorithm (Bradski, 2000) across various levels of the Differential of Gaussian (DoG) pyramid, acting as essential structural anchors. Subsequently, a 128-dimension SIFT descriptor is computed to capture the gradient landscape around each key point. The SIFT descriptor effectively represents amino acid interaction patterns, incorporating information from multiple amino acids proximal to the key point.

**Figure 1.**
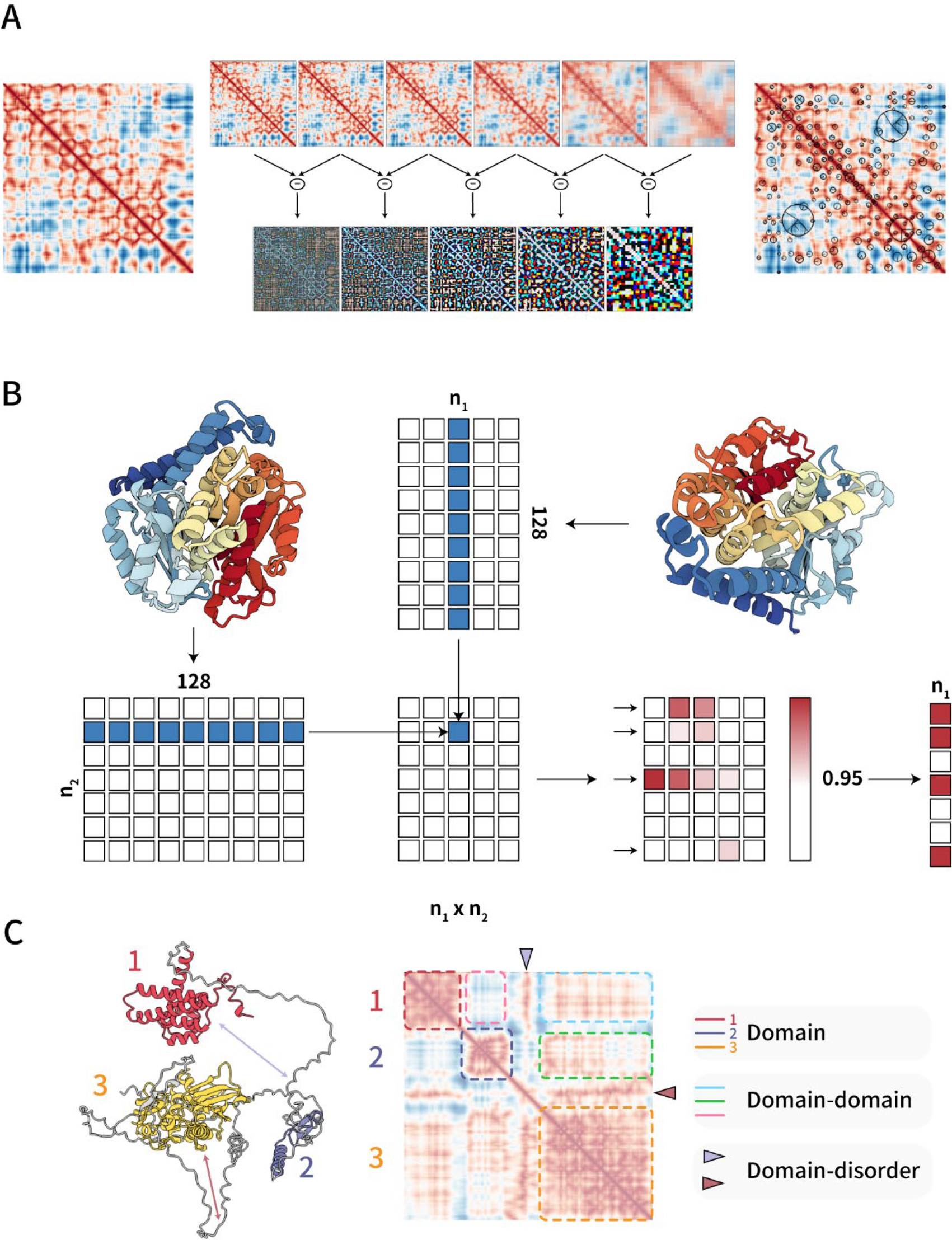
Overview of the ADAMS algorithm. A) A distance matrix is generated from the 3D coordinates of the *C*_*α*_ backbone atoms in a protein structure by calculating all-versus-all distances. (left). This matrix is then subjected to Gaussian smoothing using multiple Gaussian kernels of increasing standard deviation, creating multiple layers. The smoothed matrix is down sampled to half its size, followed by additional rounds of Gaussian smoothing, resulting in a Gaussian pyramid. (middle up) Subtracting adjacent levels of this pyramid generates a Difference of Gaussian (DoG) pyramid, highlighting edges and key features at multiple scales (middle down). A 128-dimension SIFT descriptor is generated around each key point located on the distance matrix to describe gradient changes and overall information around it. (right). B) Feature matrices are constructed from distance matrices. For each feature extracted from a distance matrix, a corresponding *i* × 128 feature matrix is created, with each row representing a standard 128-dimension SIFT descriptor. When comparing two matrices, the cosine similarity is calculated using matrix dot-products. A threshold is applied to filter out poor matches, and the number of matches and match scores are computed (see also Algorithm 1). C) An example distance matrix is provided, with domain information marked using distinct colors. Domain-domain interactions and domain-loop interactions are separately recorded in different regions of the distance matrix.

For structural comparisons, ADAMS utilizes an adapted version of the SIFT algorithm. While the conventional SIFT depends on K-nearest neighbor or direct brute-force methods to compute the Euclidean distance between SIFT vectors (Lowe, 2004), ADAMS normalizes these vectors to unit vectors and computes cosine similarity as a replacement for the distance (Fig. 1B). This adaptation augments computational efficiency, priming ADAMS for GPU acceleration. Consequently, proteins with structural similarities will present a shared ensemble of keypoints, thereby enhancing the accuracy of their structural alignment.

ADAMS provides a solution to deal with the challenges inherent in conventional protein structure alignment methods, such as Dali, CE, and TM-align (Fig. 1C). Classic alignment methodologies often grapple with the predicament while optimizing the alignment of one protein segment inadvertently compromises the alignment of another. By capturing local structure domains along the distance matrix’s diagonal, ADAMS eliminates this issue, enabling precise alignment even for proteins exhibiting flexible domains. In contrast to local structure embedding techniques like Foldseek, where domain details can get entangled with inter-domain interactions, ADAMS segregates domain information, safeguarding against potential interference from disordered or flexible regions (Fig. 1C).

### ADAMS matches Foldseek’s homologue finding performance

Protein structural comparison is crucial for locating protein homologues and deducing function, particularly when traditional sequence-based methods fall short. In a proteomics landscape where a singular taxonomy can encompass over 20,000 proteins, speed is key. Conventional tools for structural comparison, when applied to typical human proteome datasets, might take several days for completion. For perspective, in navigating the AlphaFold Database (AFDB) (David, et al., 2022) human proteome, CE took roughly five days for a singular protein analysis, while Dali lagged at an approximate 25 hours (Fig. 2A). Contrastingly, Foldseek’s prowess allowed it to traverse thousands to millions of structures within seconds. The *Caenorhabditis elegans* (*C. elegans*) Otholist 2 dataset mapped *C. elegans* genes to human homologues based on their primary amino acid sequences(Kim, et al., 2018). We benchmarked these tools against a human proteome dataset comprised of 20,696 protein structures, relying solely on structural data, utilizing Otholist 2 as the positive control. Here, Foldseek, aided by its high-efficiency prefilter, rounded off searches within an average of 3.7 seconds per search. ADAMS, in a close follow-up, finished at about 5.7 seconds in average (Fig. 2A).

**Figure 2.**
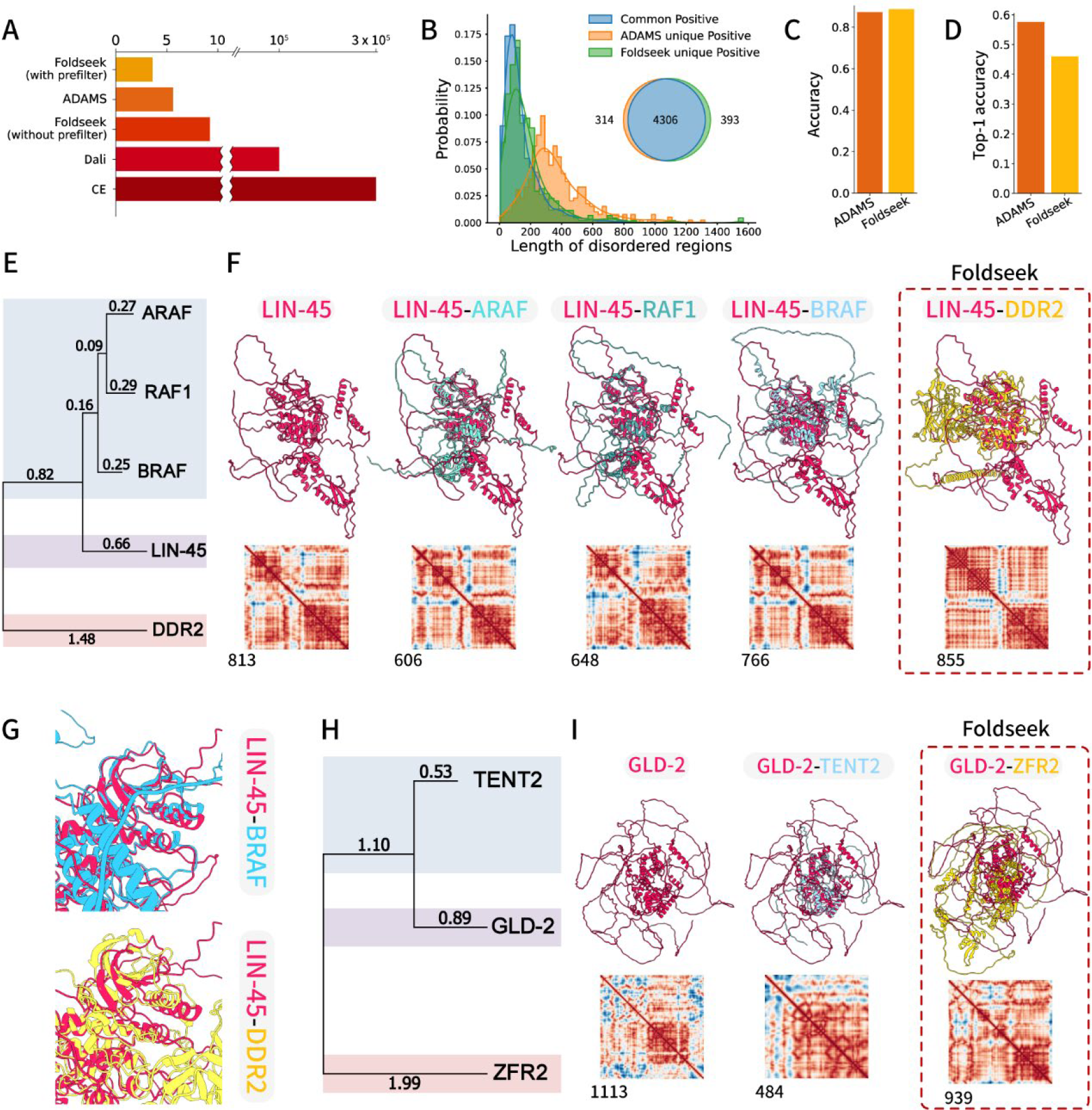
Comparison of the performance of ADAMS and Foldseek. **A)** The bar plot illustrates the average time required by ADAMS, Foldseek, Dali and CE for comparing a single protein against the entire human proteome. The bars of Foldseek and ADAMS represent the average time consumed while the bars of CE and Dali show the time spent while comparing *C. elegans* protein KSR-1 to the whole human proteome. B) The Venn diagram visually represents the count of positive matches provided by ADAMS and Foldseek when comparing *C. elegans* proteins to human proteins based on their structural features. The histogram shows the distribution of IDR length among each part of the Venn diagram. C) This bar plot demonstrates the positive rate achieved by ADAMS and Foldseek when comparing *C. elegans* proteins to human proteins by their structural characteristics. D) In this bar plot, the top-1 accuracy for the unique positive cases obtained by ADAMS and Foldseek is presented. E) A phylogenetic tree illustrates the relationship between proteins identified by ADAMS and Foldseek during the search for LIN-45. F) The results of searching for LIN-45 in the human protein database are presented for both Foldseek and ADAMS. In this comparison, ADAMS successfully identified ARAF (1), RAF1 (2), and BRAF (3) as highly similar structures. However, none of these structures were among the top 100 similar hits in Foldseek’s results. Foldseek erroneously labeled a structure as a top-ranked match, highlighted with red dashed lines, which was later confirmed to be a false positive result. G) Detailed comparison of the structure alignment of the kinase domain between LIN-45/ BRAF and LIN45/DDR2. The structures were superimposed using CE alignment. H) A phylogenetic tree depicts the relationship between proteins identified by ADAMS and Foldseek during the search for GLD-2. I) The results of searching for GLD-2 in the human protein database are provided by both Foldseek and ADAMS. ADAMS successfully identified TENT2(1). Foldseek erroneously labeled a structure as a top-ranked match, highlighted with red dashed lines, which was later confirmed to be a false positive result.

When filtering the top 100 matches, ADAMS discerned 4,620 sequence homologues with structural resemblances, closely matching Foldseek’s identification of 4,699 (Fig. 2B, C). Most of the positive results are high-ranking results (Fig. S1C). Intriguingly, ADAMS outperformed in certain niches, uncovering 314 structurally analogous proteins that Foldseek missed, underscoring its adeptness in structure alignment, most of these unique findings were containing a certain length of IDRs (Fig. 2B). Of these unique identifications by ADAMS, a notable 57.6% aligned seamlessly with sequence homologues (Fig. 2D). Conversely, Foldseek identified 393 structural homologues elusive to ADAMS, with a top-1 accuracy of about 46.1% (Fig. 2D). This comparative analysis underscores the individual strengths of both ADAMS and Foldseek: each has a distinct edge in recognizing specific structural parallels that might elude the other. Their combined capabilities offer a holistic and enriched avenue to probe into protein structures and their evolutionary interconnections.

### ADAMS filters noise from IDRs

When examining the amino acid sequence of proteins, the *C. elegans* LIN-45 protein was discerned as the counterpart of human proteins ARAF, BRAF, and RAF1 (Fig. 2E) (Kim, et al., 2018). Searching through the human proteome structures in the AFDB, ADAMS exhibited remarkable accuracy by not only detecting these three proteins but also ranking them in top three hits (Fig. 2F). In contrast, Foldseek did not fare as well, missing out on identifying any of these proteins. Moreover, Foldseek’s primary match, the kinase DDR2, appeared to be a distant relation when evaluated based on primary sequence analysis (Fig. 2E). This divergence was made evident by a distinct misalignment in Foldseek’s primary selection (Fig. 2G). These findings accentuate ADAMS’ superior capability in this specific instance.

Turning the spotlight to another *C. elegans* protein, GLD-2, ADAMS once more displayed its precision, flawlessly identifying its human counterpart, TENT2, and underscoring a notable structural affinity between them. However, Foldseek manifested a clear misalignment during this specific assessment (Fig. 2H, I). Similar consequence was also observed while comparing SMG-2 to human protein structures (Fig. S1D, E). Further, primary sequence alignment revealed that *C. elegans* KSR-1 is the homologue with two human proteins, KSR1 and KSR2 (Fig. 3A). While ADAMS recognized both KSR1 and KSR2, Foldseek only acknowledged KSR1 as the structural homologue (Fig. 3B). This is intriguing given the pronounced similarity between the two in terms of their three domains and overarching structure (Fig. 3C, D, E)

**Figure 3.**
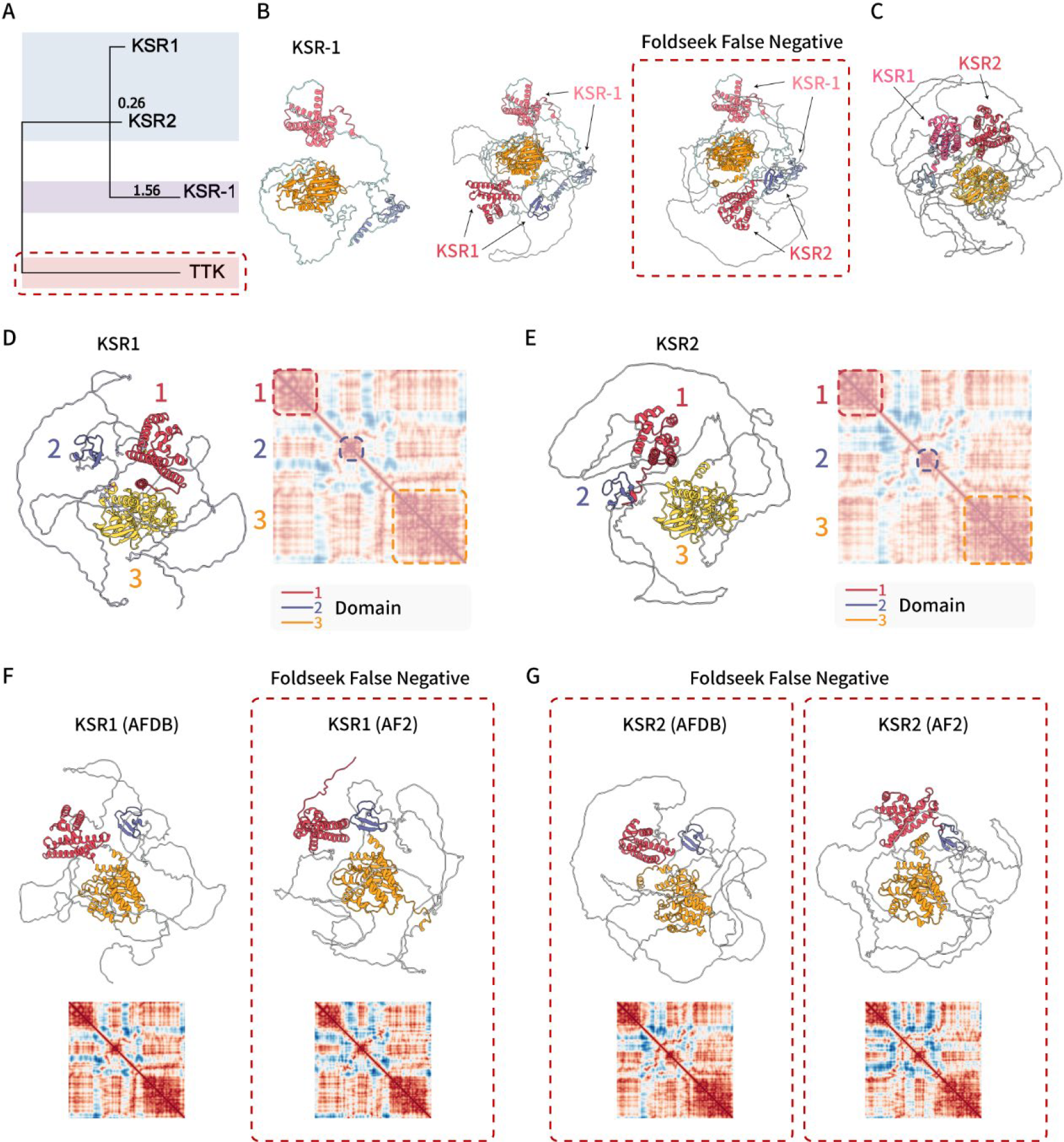
ADAMS reduces noise interactions by embedding disordered domains separately. A) A phylogenetic tree illustrates the relationship between proteins identified by ADAMS and Foldseek during the search for GLD-1. B) The results of searching for KSR-1 in the human protein database are displayed for both Foldseek and ADAMS. Red dashed lines highlight a false negative result generated by Foldseek, indicating a missed match. C) A structure alignment result showcases the structural comparison between the human proteins KSR1 and KSR2. D) Distance matrix presentation of KSR1 E) Distance matrix presentation of KSR2 F) Structural representations of KSR1 in the AlphaFold Database (AFDB) and an additional prediction from AlphaFold2 are shown. Structures within red dashed lines indicate false negative results produced by Foldseek. G) Structural representations of KSR2 in the AlphaFold Database (AFDB) and an additional prediction from AlphaFold2 are displayed. Structures within red dashed lines indicate false negative results generated by Foldseek.

We hypothesized that the observed inconsistency might stem from Foldseek’s utilization of 3Di for embedding local structural details. In situations where there are variations in flexible domains, disordered regions, or in structures that pose challenges for accurate predictions by AlphaFold2, there’s a possibility that noise information gets embedded alongside actual domain contexts. This type of noise might also be present in other coordinate-based comparison algorithms such as CE (Fig. S1F), which missed these two structures and identified two other partially similar structures as the most similar ones. This could give rise to inaccuracies in detecting structural homology, either in the form of false positives or false negatives.

To verify this hypothesis, we conducted a new prediction for the structures of KSR1 and KSR2 using ColabFold (Fig. 3F, G) (Mirdita, et al., 2022). While we did notice minor discrepancies between these newly predicted structures and the ones sourced from the AFDB, the majority of the structures were largely consistent. This fact was further substantiated by the distance matrix (Fig. 3F, G). Interestingly, when we juxtaposed these structures with other human proteome structures from the AFDB, Foldseek consistently missed detecting any version of KSR2, and the new prediction of KSR1 similarly went undetected. In contrast, ADAMS displayed unwavering consistency, successfully identifying and ranking all these structures prominently. Dali, which relies on distance matrix alignment similar to ADAMS, remained unaffected by these fluctuations and successfully identified both homologues. This underscores that the standout capability of ADAMS to adeptly manage the noise instigated by disordered regions is fundamentally linked to its specialized modus operandi. The result of overall comparisons also shows ADAMS’s unique ability in identifying proteins with long IDRs, which consist most of ADAMS unique positive cases comparing with Foldseek (Fig. 1C). By leveraging the distance matrix, ADAMS separately document domain information, including domain-domain interactions, thereby facilitating more precise protein structural comparisons.

### Limitations of ADAMS

While ADAMS consistently showcases robust sensitivity and adept noise filtration, there are scenarios where it falls short of Foldseek’s performance. For instance, in probing the structural homologues of *C. elegans* IFT-81, ADAMS missed the direct counterpart, IFT81 (Fig. 4A). This lapse can be traced back to inherent limitations of the SIFT algorithm, which struggles with structures which will produce an oversimplified distance matrix that hinders effective feature extraction. An evaluation of feature counts for proteins aptly detected by ADAMS (true positives) versus those it overlooked (false negatives) revealed that proteins from the latter category generally exhibit fewer features, corroborating our initial hypothesis (Fig. 4B). Interestingly, for certain structures like GASR-8, both ADAMS and Foldseek were unable to pinpoint the direct counterpart GAS8 within their top 100 hits, despite the foremost hit showcasing a commensurate structure and distance matrix (Fig. 4C). This underscores the notion that for some proteins, mere structural comparison might be insufficient, necessitating supplementary sequence alignment.

**Figure 4.**
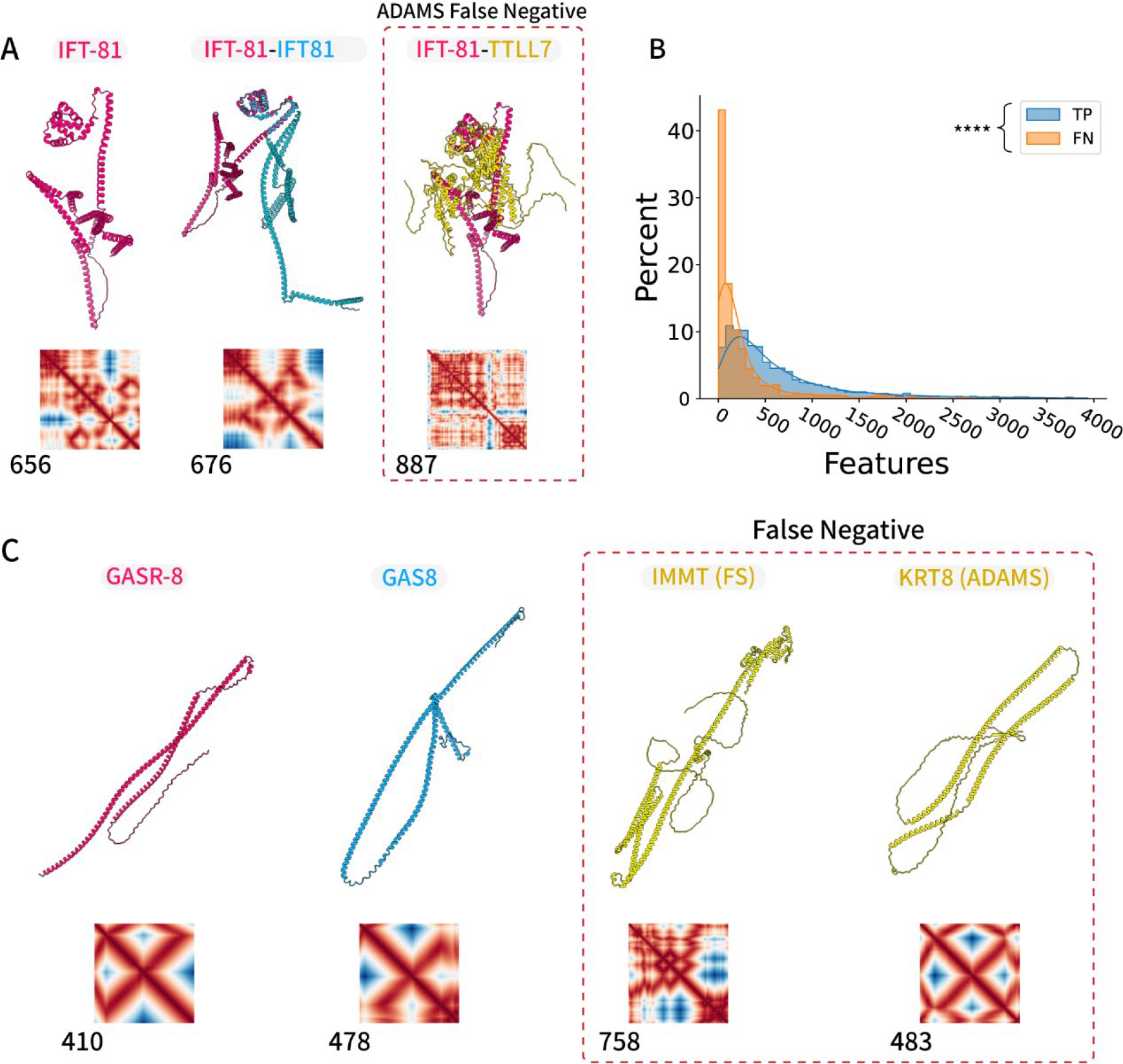
False negative results of ADAMS or Foldseek. A) The results of searching for IFT-81 in the human protein database are presented for both Foldseek and ADAMS. Notably, red dashed lines highlights a false positive result produced by Foldseek. B) Distribution of feature count in ADAMS false negative proteins and true positive proteins. **** p<0.000, Welch’s t-test. C) The results of searching for GASR-8 in the human protein database are displayed for both Foldseek and ADAMS. In this case, red dashed lines indicate instances where both Foldseek and ADAMS generated false positive results.

## Discussion

ADAMS introduces a sophisticated method for navigating the complex domains of protein structures. Echoing Dali’s foundational principle of distance matrix comparison, it leverages computer vision algorithms to dramatically slash computational demands by several orders of magnitude, allowing ADAMS to adeptly address challenges that traditionally stymie conventional protein structure comparison tools. ADAMS attains an overall efficacy and precision with Foldseek, matched by a comparable rapidity. Notably, ADAMS mitigates some of Foldseek’s shortcomings in analyzing proteins with flexible structures, whereas Foldseek counterbalances the issue of ADAMS’ occasional paucity of SIFT features necessary for structural alignment. The correlation between feature count and accuracy in ADAMS’ detection underscores the algorithm’s inherent dependency on the richness of structural features. Proteins with a simpler structural profile, reflected in a lower feature count, potentially lie outside the optimal detection range of ADAMS. This observation aligns with the classical trade-off often encountered in bioinformatics: that between sensitivity and specificity.

Structure embedding, a method increasingly embraced by contemporary structure comparison tools, transforms the complexity of 3D coordinate comparison into streamlined sequence or vector-based evaluations. Pioneering platforms like CLE (Wang and Zheng, 2008), 3D-Blast (Yang and Tung, 2006), and Foldseek utilize distinct structure alphabets and substitution matrices. In contrast, emergent tools, especially those harnessing deep learning (Greener and Jamali, 2022), gravitate towards embedding holistic protein structural information into vectors, thereby amplifying comparison efficiency. However, the intrinsic limitation of structure embedding stems from its potential imprecision, given its departure from the exhaustive n*n distance matrix representing protein structures. In this embedding transition, information is often compacted, typically along the matrix’s diagonal or even into more concise sequences. This compression might inadvertently integrate extraneous or irrelevant data, compromising the fidelity of the original information and thereby muddling the accurate depiction of structural nuances.

The deployment of distance matrices for structural analysis, a concept originally advanced by Dali, offers a potent modality to discern complex interaction patterns among protein residues, furnishing a detailed representation unencumbered by data reduction. This methodology’s adaptability is notably advanced through the ADAMS algorithm, which incorporates this approach with broad applicational scope. Such adaptability is invaluable for examining proteins with flexible domains. In the current field of structural biology, replete with vast databases brimming with Artificial Intelligence (AI) generated structures, the recognition of potential influences - ranging from AI model parameter variances to model version upgrades, and even hardware precision discrepancies - is essential. Distance matrices stand out as a robust tool to maintain data fidelity and facilitate structural comparisons.

Furthermore, the applicability of distance matrix-centric approaches encompasses evaluations of protein complexes and emergent Cryo-Electron Microscopy (Cryo-EM) structures. In these scenarios, traditional structure embedding techniques may struggle due to their sensitivity to structural variations and their limited utility in low-resolution Cryo-EM structures (Sorzano, et al., 2017), which often exhibit inherent complexities arising from undesirable interactions. In contrast, leveraging distance matrices offers a more resilient search approach for these intricate structures, underscoring their potential in advancing structural biology.

While both ADAMS and Foldseek are tailored for systematic protein structure comparisons, they overlook the direct homologue of GASR-8 (Fig. 4C), highlighting that exclusive reliance on structural data can sometimes be limiting. These instances emphasize that no individual strategy stands as a definitive solution within structural bioinformatics. Rather, an integrated approach, harnessing the virtues of various algorithms, seems to be a promising avenue. Merging structural analyses with sequence alignment and other bioinformatics methods may allow researchers to surmount prevalent challenges, fostering a richer and more intricate understanding of protein structures and their functional implications.

## Materials and Methods

### Overview

ADAMS revolutionizes protein structure comparison by integrating Distance Matrix Alignment (Dali) with the SIFT algorithm. In this advanced approach, cosine similarity is employed to assess the distance between each SIFT descriptor, optimizing for swift and precise comparisons between protein structures. Notably, ADAMS leverages GPU acceleration, ensuring rapid and detailed distance matrix alignment. Furthermore, it stands out as the fastest tool for this purpose, with the versatility to run on a wide array of GPU devices, all while keeping GPU memory consumption low (typically 500-2000MB per comparison) and allowing for multi-process capabilities. All the GPU acceleration process in this article were done with a single NVIDIA RTX2080Ti (11 GB) GPU.

### SIFT algorithm generating 128-dimension key point descriptions from distance matrix

The Basic SIFT algorithm closely follows the methodology described in the original article. It involves several key steps:

1. Normalization of Distance Matrix: The first step involves normalizing the distance matrix to a 0-255 scale image: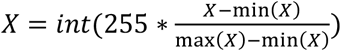.
2. Scale Space Extreme Value Detection: This step identifies extreme values within the scale space to pinpoint potential keypoints.
3. Key Localization: Once extreme values are detected, keypoint localization is performed to precisely determine the location of keypoints in the image.
4. Orientation Determination: After localization, the algorithm calculates the orientation of each keypoint to account for the direction of local features.
5. Key Point Description: In this crucial step, a 128-dimensional vector is generated to describe the gradient information around each keypoint. This vector takes into account the position, scale, and orientation of the keypoint.
6. Descriptor Normalization: To mitigate the impact of non-linear illumination changes, the descriptors are normalized using the as: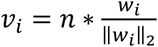. Here, *v* _*i*_ represents the normalized vector and *w*_*i*_ represents the original descriptor.

In our approach, we leverage the well-established and robust OpenCV-SIFT algorithm, which has consistently demonstrated its effectiveness and reliability in a wide range of applications.

### Advanced key point matching algorithm

In traditional SIFT keypoint matching algorithms, the process of determining the best matching keypoints often involves the use of the K-nearest neighbor (KNN) or Brute-force matching approach. This method relies on the Euclidean distance to assess the similarity between invariant descriptor vectors of keypoints. However, in relatively small-scale images, obtaining a sufficient number of keypoints for KNN modeling can be challenging, and performing Euclidean distance calculations for searching through billions of keypoints can be computationally intensive.

In the context of protein structure comparison, the characteristics of the data differ from typical camera pictures for which SIFT was originally designed. Specifically, the distance matrix in this context tends to have minimal noise, and the orientation and scale levels are relatively consistent across different proteins. As a result, an alternative approach is employed, substituting the Euclidean distance calculation with cosine similarity for a faster, albeit slightly less accurate, evaluation of the similarity between descriptor vectors. This strategy takes advantage of GPU acceleration for efficient matrix multiplication, significantly improving the speed of the process. (Algorithm 1)

### Z-score

ADAMS utilize the sum up z-score, which added up the z-score of the number of matched points of which the cosine similarity is beyond the threshold and the z-score of the match score (Algorithm 1) to rank the results.

### Dali

We utilized the standalone DaliLite.v5 for our analysis. The input files were formatted as DAT files using Dali’s import.pl. This conversion resulted in 20,566 valid structures from an initial count of 20,696 within the human proteome dataset. Following this formatting process, we computed the protein alignment utilizing Dali’s structural alignment algorithm. The results were sorted by Dali’s z-score in descending order.

DaliLite.v5/bin/dali.pl --query query.list --db id.list --dat1 match/ --dat2 DaliDatabase/ --clean

### CE

We used the same CE alignment module used in the Foldseek article (van Kempen, et al., 2023), which can be downloaded from https://github.com/steineggerlab/foldseek-analysis.

### Protein structure prediction with ColabFold and AlphaFold2

The tertiary structures of full-length KSR1 (1-923) and KSR2 (1-950) were predicted using ColabFold: AlphaFold2 using MMseqs2 (https://colab.research.google.com/github/sokrypton/ColabFold/blob/main/AlphaFold2.ipynb). The structures with highest plDDT values were selected for further analysis.

### Statistical analysis

Evolutionary distance analysis was performed by MEGA7 (version 11.0.13) software (Kumar, et al., 2016). Protein structures and superimposing were performed and displayed with UCSF Chimera, developed by the Resource for Biocomputing, Visualization, and Informatics at the University of California, San Francisco. The length and percentage of IDRs shown in this article were identified by DSSP module provided by Biopython (1.81) (Cock, et al., 2009) and the DSSP (2.1.0) (Touw, et al., 2015) program. The length of each protein was obtained from protein structure models from AFDB database. Welch’s t-test was employed in Fig. 4B to compare the number of features using scipy.stats.ttest_ind() in Python (3.11.5)

## Data availability

All the protein structure data used in this article can be downloaded from AlphaFold Protein Structure Database. Human protein structure is available from https://ftp.ebi.ac.uk/pub/databases/alphafold/latest/UP000005640_9606_HUMAN_v4.tar, *C. elegans* protein structure is available from https://ftp.ebi.ac.uk/pub/databases/alphafold/latest/UP000001940_6239_CAEEL_v4.tar. Data of ortholist2 can be obtained from http://ortholist.shaye-lab.org/send_master.

## Code availability

ADAMS can be download and used as a python package from Python Package Index (PyPI): [adams · PyPI](https://pypi.org/project/adams/). Source code and other materials are available from young55775/ADAMS-developing (github.com). An online server is available: https://cryonet.ai/bseek.

### Algorithm 1: Key point matching by cosine similarity

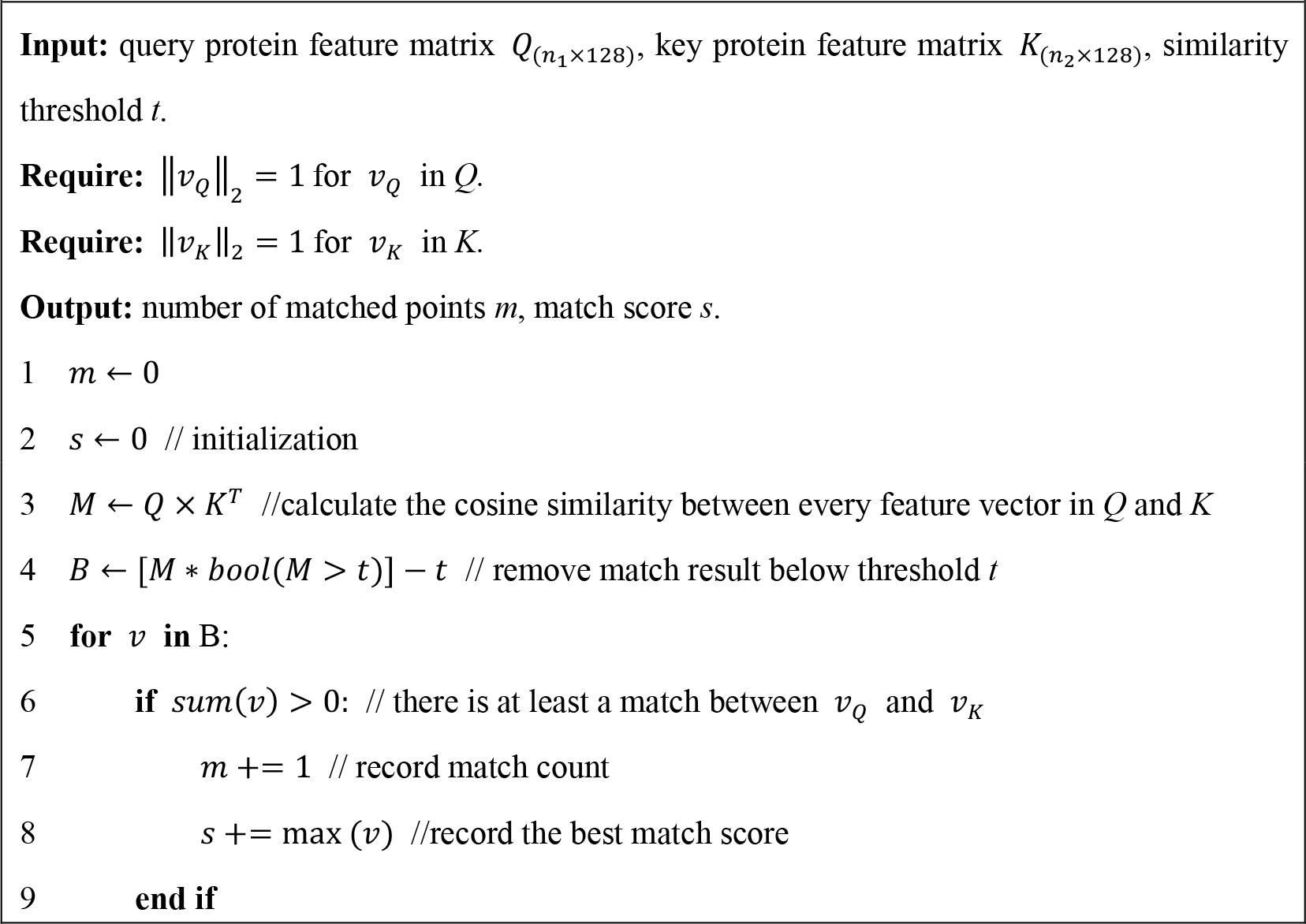
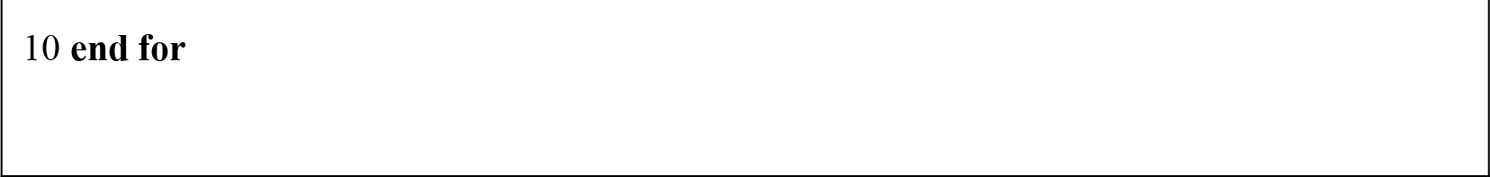

## Acknowledgements

We thank Drs. K. Xu and C. Zhang for discussion. This work was supported by the National Natural Science Foundation of China Grants (31991190, 31730052, 31861143042, 31671444, and 31871352) and National Key R&D Program of China Grants (2019YFA0508401 and 2017YFA0102900).

## Author Contributions

G.Y., Y. W., and G.O. designed the project. G.Y. and Y. W. performed the analysis. G.Y. and G.O. wrote the paper.

