## Supplementary Figure 1 for "Utilizing the Scale-Invariant Feature Transform Algorithm to Align Distance Matrices Facilitates Systematic Protein Structure Comparison"

**Figure S1**
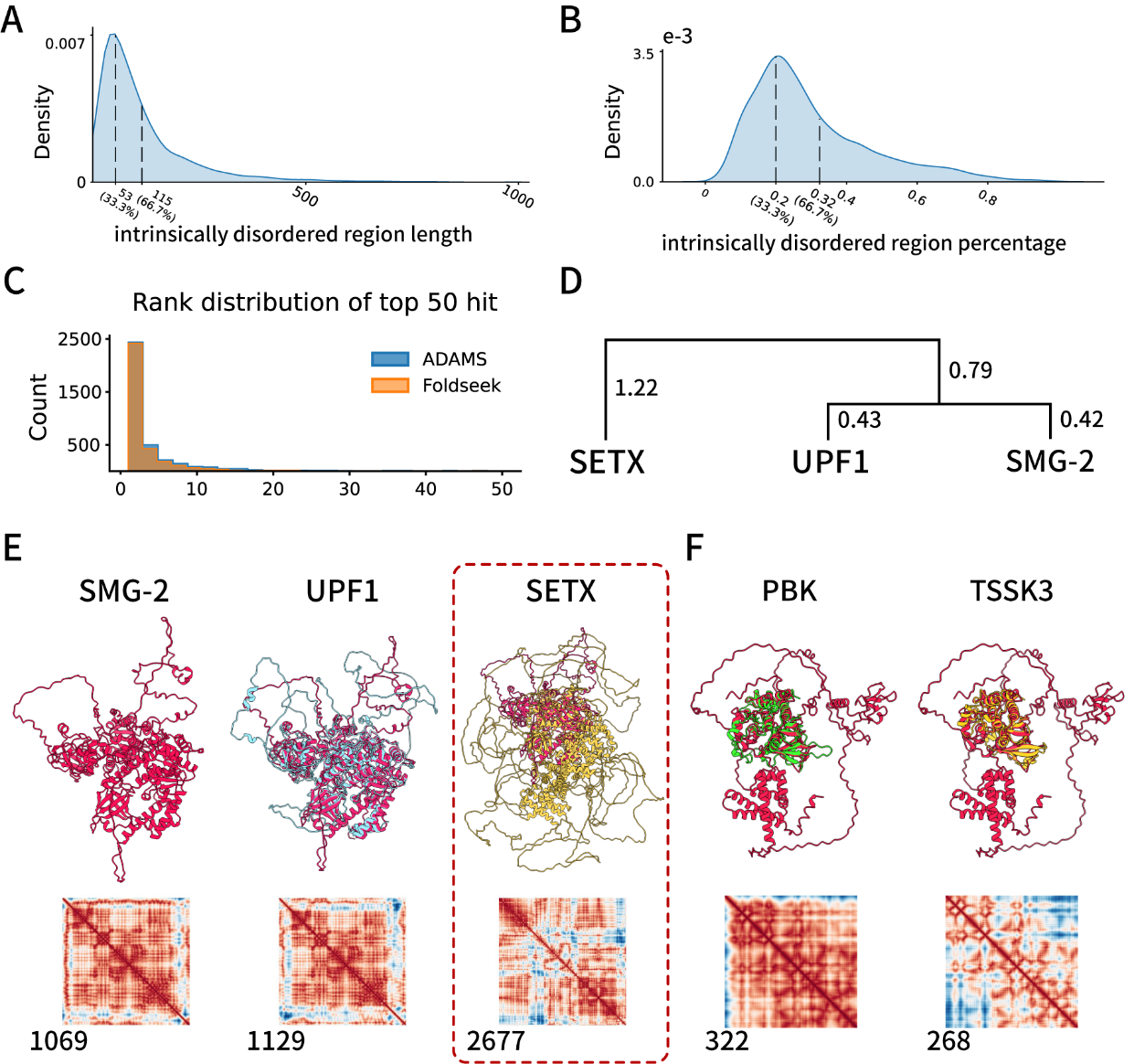

**Figure S1. ADAMS filter noise from IDR rich proteins**

A) Length distribution of intrinsically disordered regions in human proteins.
B) Percentage of intrinsically disordered regions in human proteins.
C) Distribution of positive hit rank of the positive cases when the positive case was in top 50 hit of both methods.
D) A phylogenetic tree illustrates the relationship between proteins identified by ADAMS and Foldseek during the search for SMG2.
E) The results of searching for SMG-2 in the human protein database are provided by both Foldseek and ADAMS. ADAMS successfully identified UPF1(1). Foldseek erroneously labeled a structure as a top-ranked match, highlighted with red dashed lines, which was later confirmed to be a false positive result.
F) The top rank results of searching for KSR-1 in the human protein database provided by CE.
